# First detection and characterization of Alongshan virus in *Ixodes ricinus* ticks from Italy, 2021-2022

**DOI:** 10.64898/2026.06.13.732040

**Authors:** Silvia Fabi, Mariachiara Vardeu, Alex Martini, Elisa Franchin, Elisabetta Valente, Fabrizio Montarsi, Graziana Da Rold, Federica Obber, Chiara Agostini, Arianna Breda, Claudia Del Vecchio, Ignazio Castagliuolo, Enrico Lavezzo, Cristiano Salata

## Abstract

Alongshan virus (ALSV) is an emerging tick-borne segmented RNA virus belonging to the Jingmenvirus group and has been reported in humans, ticks, and vertebrates across Asia and Europe. Despite its potential public health relevance, its distribution and genetic diversity remain poorly characterized in several European regions where tick-borne pathogens are endemic. In this study, we developed a specific TaqMan-based real-time RT-PCR assay targeting a conserved region of ALSV segment 2 and used it to investigate the presence of ALSV RNA in *Ixodes ricinus* ticks collected in northeastern Italy. The assay showed high linearity over a broad dynamic range and no cross-reactivity with related flaviviruses. A total of 212 archival tick samples collected between March 2021 and November 2022 were screened, and 28 samples (13.2%) tested positive for ALSV RNA. Positive ticks were detected in the provinces of Belluno and Vicenza and included individual adult males and nymph pools. A subset of positive samples was further characterized by nested PCR and Sanger sequencing of all four genomic segments. Phylogenetic analyses showed that Italian ALSV sequences clustered within the broader European ALSV diversity and were closely related to strains from Central and Northern Europe, without forming a distinct country-specific lineage. Sequence comparisons suggested purifying selection and revealed differences in predicted structural proteins between European and Chinese strains. These findings provide the first molecular evidence of ALSV circulation in Italy and support further studies to clarify its epidemiology, host range, genetic diversity, and potential clinical relevance.

**IMPORTANCE:** Alongshan virus (ALSV) is an emerging tick-borne virus identified in febrile patients in China and subsequently detected in ticks in Russian Federation and several European countries. Although severe disease has not yet been reported in humans, surveillance and elucidation of the virus distribution are essential to assess its pathogenicity and potential public health impact. We developed a specific real-time RT-PCR protocol and detected ALSV in *Ixodes ricinus* ticks collected in northeastern Italy. Sequence analyses suggested multiple introductions and revealed differences in structural proteins between European and Chinese strains, suggesting potential adaptation and differences in pathogenicity. Since the clinical signs of ALSV infection in humans may overlap with those of tick-borne encephalitis (TBE), differential diagnostic procedures should be developed to improve patient management, particularly in TBE-endemic regions such as northeastern of Italy.

## INTRODUCTION

The Alongshan virus (ALSV) is an emerging pathogen of public health concern, first identified in a febrile woman in China in 2017 (1). To date, this novel virus has been detected in approximately 90 Chinese patients presenting with symptoms including fever, headache, skin rash, arthralgia, myalgia, depression, and coma. Notably, no long-term sequelae or lethality have been reported (2). Molecular characterization reveals that ALSV possesses a segmented, positive-sense, single-stranded RNA genome (1). It belongs to the recently discovered Jingmenvirus (JMV) group which comprises RNA viruses with genomes consisting of four or five segments of positive-sense, single-stranded RNA (3, 4).

The ALSV genome comprises four segments of positive sense RNA, each featuring a 5’ cap and a 3’ polyA tract. Segments 1 and 3 show homologies to NS3 and NS5 open reading frames (ORFs) of classical flavivirus, while segments 2 and 4 are unique to the JMV group, lacking homology with other known sequences. Due to their partial homology with flaviviruses, ALSV and all JMV group members are classified within the *Flaviviridae* family as unclassified flaviviruses (4).

Although primarily considered a tick-borne virus, ALSV has been detected in various hosts including mosquitos, deer, sheep, and cattle (4). Since its discovery, ALSV has been identified in China, the Russian Federation, and several European countries. Specifically, it has been found in *Ixodes ricinus* ticks collected in Finland, France, Switzerland, Poland, Serbia, and Austria (5–10), in both *I. ricinus* and *Dermacentor reticulatus* in Germany, with evidence of circulations in vertebrates, and in *I. hexagonus* in United Kingdom (11).

A recent study by Janshoff and colleagues reported the detection of anti-ALSV antibodies in commercial serum pools from horses in the Americas, Europe, and Oceania, suggesting widespread distribution and high circulation of the virus (12).

*Ixodes ricinus* ticks are prevalent throughout central and northern Italy, with the northeastern region being an endemic hotspot for tick-borne pathogens, including the tick-borne encephalitis virus (TBEV) (13, 14). The clinical presentation of ALSV infection often resembles that of tick-borne encephalitis (TBE) (1), highlighting the crucial need to confirm ALSV presence in the region. This information is essential for enhancing surveillance strategies, improving differential diagnosis protocols, and optimizing patient management.

Here, we report the detection and molecular characterization of ALSV RNA in *I. ricinus* ticks collected in northeastern Italy, a TBE-endemic area, using a newly developed specific real-time RT-PCR assay. Our findings provide the first molecular evidence of ALSV circulation in Italy and highlight the need of further studies to assess its distribution, genetic diversity, and potential impact on human and animal health in TBE-endemic regions of Europe.

## RESULTS AND DISCUSSION

We analyzed 212 archival samples derived from *I. ricinus* ticks collected in northeastern Italy between March 2021 and November 2022 (13) to investigate the presence of ALSV. The dataset included individual adult ticks and pools of up to ten larvae or nymphs. Overall, ALSV RNA was detected in 28 samples (13.2%), all originating from the provinces of Belluno and Vicenza. Positive samples included four individual males and multiple nymph pools (Table). The detection of ALSV RNA in samples collected over two consecutive years suggests that the virus is not sporadically introduced but is likely established within local tick populations in northeastern Italy.

Detection was performed using a newly developed TaqMan-based real-time RT-PCR assay targeting a conserved region of genomic segment 2. The assay showed high linearity (R^2^ = 0.9992) and no cross-reactivity with related flaviviruses tested, namely dengue virus serotypes 1 and 2, West Nile virus lineages 1 and 2, yellow fever virus, tick-borne encephalitis virus, and Zika virus. These results confirm the specificity and robustness of the assay for ALSV detection in tick-derived samples in ecological settings where multiple flaviviruses co-circulate.

A subset of 15 positive samples was further characterised by nested PCR targeting all four ALSV genomic segments, followed by Sanger sequencing. Overlapping amplicons of approximately 500 bp were generated (Table A1) and used to reconstruct viral genomic segments. Consensus sequences were generated by mapping Sanger reads to a curated ALSV reference database and deposited in GenBank (Table).

Phylogenetic analysis based on genomic segment 4 showed that all Italian sequences clustered within the previously described European ALSV diversity, without forming a distinct country-specific lineage (Figure 1). Italian strains were closely related to sequences from Central and Northern Europe, particularly from Switzerland and Germany, and were distributed across multiple clusters, some of which were moderately to highly supported. Similar clustering patterns were observed for genomic segments 1–3 (Figures A1–A3), although minor differences should be interpreted cautiously due to the use of partial genomic regions.

**Figure 1.**
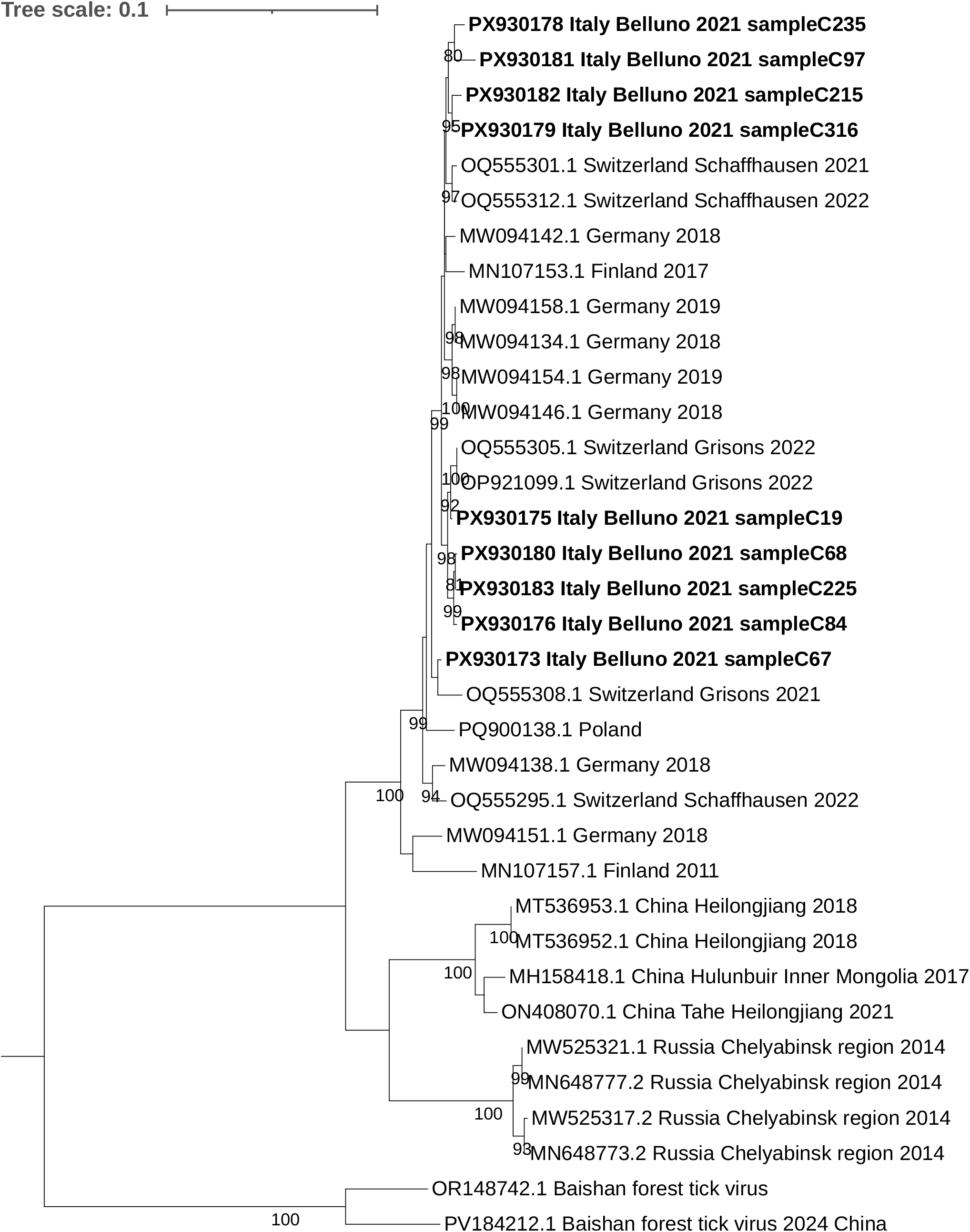
Phylogenetic tree of ALSV segment 4 sequences, including strains detected in *Ixodes ricinus* ticks, northeastern Italy, 2021–2022. Italian sequences (bold) cluster within the European ALSV diversity and are closely related to strains from Switzerland and Germany. The tree was inferred using the Maximum Likelihood method with the Tamura–Nei model. Bootstrap values (>75%, 500 replicates) are shown. Scale bar indicates nucleotide substitutions per site.

Overall, these findings indicate that ALSV circulating in northeastern Italy is part of a broader European viral population. The absence of a distinct Italian lineage, together with the distribution of sequences across multiple clusters, suggests a scenario of multiple introductions, local circulation, or both, rather than the emergence of a geographically restricted lineage.

To further investigate the evolutionary dynamics of ALSV, codon-based selection analyses were performed on all coding regions using the Datamonkey platform (15). Sequences containing more than 20% ambiguous nucleotides were excluded from downstream analyses to minimize the impact of missing data. Across all genomic segments, global dN/dS ratios estimated using SLAC were consistently below 1, ranging from 0.025 to 0.286, indicating pervasive purifying selection acting on ALSV coding regions. The strongest selective constraints were observed in segment 1 (dN/dS = 0.025) and segment 4 ORF2 (dN/dS = 0.041), whereas segments 2 ORF1 and ORF2 showed comparatively higher values (dN/dS = 0.233 and 0.286, respectively), suggesting a more relaxed selective pressure in these regions.

Site-specific analyses using FEL identified a large number of codons under significant purifying selection and no sites under pervasive diversifying selection, whereas MEME detected a limited number of codons under episodic diversifying selection.

These findings are consistent with the predominance of synonymous over non-synonymous substitutions observed in sequence comparisons. Several amino acid substitutions were shared among Italian and other European sequences, indicating substantial conservation within the European ALSV population. Conversely, only a limited number of mutations were unique to individual Italian sequences (Figure 2, Table A2, and Figures A4–A6). Overall, these results indicate that ALSV evolution is primarily driven by purifying selection, with only limited evidence of episodic adaptive changes.

**Figure 2.**
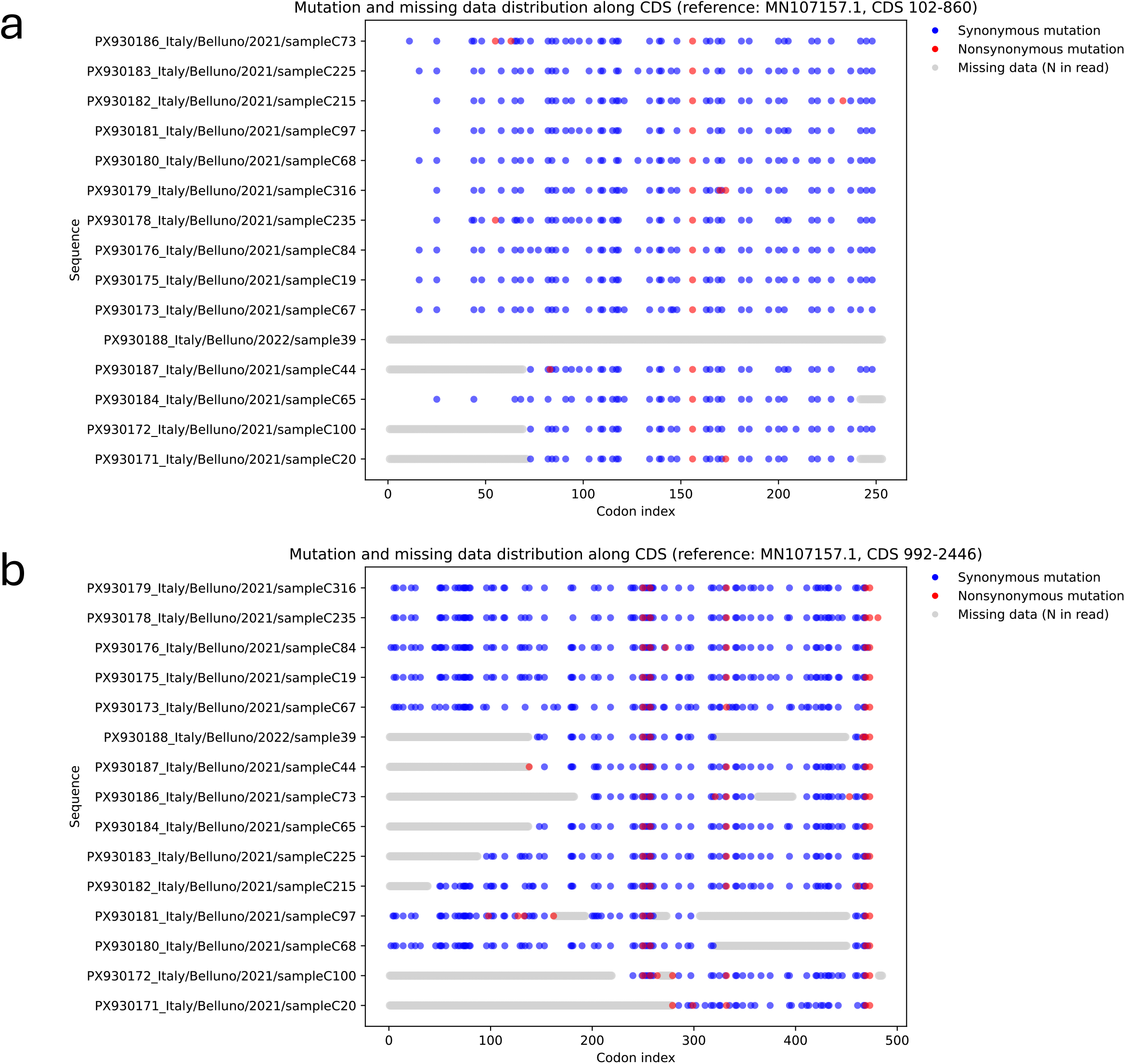
Summary of all mutations detected in segment 4 of Italian ALSV strains relative to the Finnish European reference sequence. Synonymous and non-synonymous mutations are reported as blue and red dots, respectively.

Comparative analysis of structural protein-coding segments, namely segments 2 and 4, revealed differences between European strains and previously reported Chinese ALSV strains. In particular, the predicted start codons of the VP1b (glycoprotein) and VP3 (membrane protein) were located downstream in European strains, including Italian sequences, compared with Chinese strains. These shifts, corresponding to 207 and 162 nucleotides, respectively, result in shorter predicted proteins. Interestingly, these structural protein-coding regions also displayed relatively higher dN/dS values compared with replication-associated segments, suggesting that they may be subject to more relaxed evolutionary constraints. Although the biological significance of these differences remains unknown, they may reflect adaptation to different ecological or host environments (16).

Taken together, the phylogenetic and evolutionary analyses indicate that ALSV circulating in northeastern Italy is part of a broader European viral population evolving under strong purifying selection. The detection of multiple positive samples across consecutive years supports stable local circulation, while the absence of a distinct Italian lineage suggests ongoing viral exchange across European regions. Given the overlap in ecological niches and clinical presentation with tick-borne encephalitis virus, continued surveillance and improved diagnostic differentiation are warranted to better assess the epidemiological and clinical relevance of ALSV in endemic areas.

## MATERIALS AND METHODS

### TaqMan primers and probe design

All complete ALSV genome sequences available in GenBank (accessed on 09/2025) were retrieved and aligned to identify conserved regions across the four genomic segments. A conserved region within segment 2 was selected as the target for the real-time RT-PCR assay development. Segment 2 was selected due to the absence of homology with other viruses. The primers ALSVFw-CACATCACGGGAGGTATCG and ALSVRev-TGACCACAAGCACAGTTGGA, together with the TaqMan probe ALSVPr-CGGATYATCTCATGGGCGGTCG, were designed using Primer-BLAST 2.17.0 (NCBI) and synthesized by Biosense s.r.l. (Milano, Italy).

### Preparation and quantification of positive control

The positive control for the real-time PCR assay, containing the target sequence selected in this study, was produced by annealing two complementary 92-bp oligonucleotide strands derived from segment 2 of ALSV. The resulting double-stranded oligonucleotide was cloned into the pJET1.2/blunt vector (Thermo Fisher Scientific, Waltham, MA, USA) according to the manufacturer’s instructions and subsequently propagated in *Escherichia coli* DH5α cells. The obtained pJET-ALSV plasmid was subjected to Sanger sequencing to confirm the correct insert sequence.

Droplet digital PCR (ddPCR) was used for the absolute quantification of the pJET-ALSV plasmid using the QX200™ Droplet Digital PCR System (Bio-Rad Laboratories, Berkeley, CA, USA), as previously described (17, 18). The reaction mixture contained 11 µL of ddPCR™ Supermix (Bio-Rad), 3 µL of primer–probe mix, 3 µL of nuclease-free water, and 5 µL of sample. The primer– probe mix was prepared by combining 22.5 µL of forward primer (100 µM), 22.5 µL of reverse primer (100 µM), 5 µL of probe (100 µM), and 250 µL of nuclease-free water. Droplets were generated, transferred to a 96-well plate, and amplified under the following cycling conditions: an initial denaturation step at 95 °C for 10 min, followed by 45 cycles at 94 °C for 30 s and 60 °C for 1 min, and a final 10-min step before droplet reading. Quantification was performed using QuantaSoft™ Software v1.7, which provided the absolute copy number per microliter of the initial sample.

### Real-time PCR

The assay was based on a TaqMan™ hydrolysis probe and performed using the designed primers and probe. The 25-µL reaction mixture contained 12.5 µL of TaqMan™ Universal PCR Master Mix 2X (Thermo Fisher Scientific), 2 µL of primer–probe mix, 10 µL of pJET-ALSV plasmid, and PCR-grade nuclease-free water. The primer–probe mix was prepared by combining

22.5 µL of forward primer (100 µM), 22.5 µL of reverse primer (100 µM), 5 µL of probe (100 µM), and 250 µL of nuclease-free water. Cycling conditions were as follows: 2 min at 50 °C and 10 min at 95 °C for DNA polymerase activation, followed by 40 cycles of 95 °C for 10 s and 60 °C for 1 min. Reactions were performed on a QuantStudio™ 7 Pro System (Thermo Fisher Scientific). The analytical performance of the assay was evaluated using serial dilutions of the pJET-ALSV plasmid quantified by droplet digital PCR. The assay showed high linearity over a broad dynamic range (2.98 × 10^14 to 2.98 × 10^5 copies/µL), with a coefficient of determination (R^2^) of 0.9992. The limit of detection was estimated at 2.52 × 10^5 copies/µL.

### Total nucleic acid extraction and One-step real-time reverse-transcriptase (RT)-PCR

Total nucleic acids were purified from 200 µL of tick homogenate with the MagNA Pure 96 System (Roche Applied Sciences, Basel, Switzerland), following the manufacturer’s protocol.

The recovered nucleic acid was tested for the presence of ALSV by one-step real-time RT-PCR using the AgPath-ID™ One-Step RT-PCR Kit (Thermo Fisher Scientific). Each reaction was carried out in a final volume of 25 µL, including 12.5 µL of 2X RT-PCR Buffer, 1 µL of 25X RT-PCR Enzyme Mix, 3 µL of primers-probe mix, 5 µL of nucleic acids extract, and PCR-grade nuclease-free water. The primer-probe mix was obtained by adding 22.5 μL of forward primer (100 µM), 22.5 μL of reverse primer (100 µM), 5 µL of probe (100 µM), and 250 μL of nuclease-free water. The thermal profile consisted of reverse transcription at 50 °C for 10 min, followed by RT inactivation and polymerase activation at 95 °C for 10 min, and 40 amplification cycles of 95 °C for 10 s and 60 °C for 1 min. The pJET-ALSV plasmid was used as positive control.

### Reverse transcription

The samples that tested positive to ALSV in the one step real-time RT-PCR, were subjected to reverse transcription, using HiScript III RT SuperMix for qPCR (+gDNA wiper) Kit (Vazyme, Nanjing, China). Each mixture containing 10 µL of PCR grade water, 4 µL of gDNA wiper mix and 2 µL of total nucleic acids extract was subjected to a step at 42 °C for 2 min to eliminate contaminating genomic DNA present in the sample. Then, 4 µL of HiScript III RT SuperMix were added to each mixture before the following incubations: 37 °C for 15 min for the reverse transcription and 85 °C for 5 s for the enzyme inactivation.

### End-Point PCR and sequencing

ALSV cDNAs were amplified adopting a nested-PCR approach, employing the AmpliTaq Gold™ DNA Polymerase with Gold Buffer and MgCl_2_ (Thermo Fisher Scientific). For each sample, 4 µL of cDNA were used as a template in a first conventional PCR, employing for each of the four genomic ALSV segments an external pair of primers (seg1 FW1-RV7; seg2 FW1-RV7; seg3 FW1-RV5; seg4 FW1-RV6, Table A1). Then, 1 µL of the first PCR was subjected to a second PCR, using internal primer pairs (Table A1).

Each PCR mixture contained 5 µL of 10x Gold PCR buffer, 5 µL of 2 mM dNTPs mix, 3 µL of MgCl_2_ 25 mM, 1 µL of each primer (10 µM), 0,25 µL of AmpliTaq Gold™ DNA Polymerase 1000 U Kit, cDNA (4 µL in the first round PCR and 1 µL in the nested PCR) and PCR grade water up to the final reaction volume of 50 µL. PCR cycling conditions were: 1 cycle of denaturation at 95 °C per 10 min, and 40 cycles of amplification at 95 °C per 30 s, 52-55 °C per 30 s, and 72 °C per 3 min followed by and additional extension step at 72 °C per 7 min. ALSV-positive samples PCR products were purified using PureLinkTM Quick Gel Extraction and PCR Purification Combo Kit (Thermo Fisher Scientific) and were submitted for Sanger sequencing using Mix2Seq Kit (Eurofins Genomics, Ebersberg, Germany).

### Sequence analysis and consensus generation

Consensus sequences were generated by mapping Sanger reads against a curated database containing all available ALSV sequences from NCBI using BWA-MEM v0.7.17-r1188 (19). For each sample, the reference sequence with the highest number of mapped reads was selected as the initial template. Reads were then realigned to the selected reference to improve mapping accuracy.

Consensus sequences were generated using a majority-rule approach at each genomic position. Positions without coverage were assigned as “N”.

### Phylogenetic analyses

Phylogenetic analyses were performed using MEGA version 12 (20). Nucleotide sequences corresponding to each ALSV genomic segment were aligned with reference sequences retrieved from GenBank. Phylogenetic trees were inferred using the Maximum Likelihood method based on the Tamura–Nei substitution model. The robustness of the inferred tree topology was assessed by bootstrap analysis with 500 replicates, and only values above 75% were considered significant.

### Selection analyses

Codon-based selection analyses were performed using the Datamonkey web server (http://www.datamonkey.org), which implements the HyPhy software package. Coding sequences corresponding to each ALSV genomic segment were aligned at the nucleotide level and manually inspected to ensure correct codon alignment. Positions containing gaps or ambiguous nucleotides were retained only if they did not disrupt the reading frame. Sequences containing more than 20% ambiguous nucleotides (N) were excluded from downstream analyses.

Maximum-likelihood phylogenetic trees were inferred automatically by the Datamonkey server for each alignment. Selection pressures were assessed using the Single-Likelihood Ancestor Counting (SLAC), Fixed Effects Likelihood (FEL), and Mixed Effects Model of Evolution (MEME) methods. SLAC was used to estimate global dN/dS ratios, whereas FEL and MEME were used to identify codon sites under pervasive and episodic selection, respectively. Statistical significance was assessed using a p-value threshold of 0.05.

## ACKNOWLEDGMENTS

This research was supported by EU funding within the MUR PNRR Extended Partnership initiative on Emerging Infectious Diseases (Project no. PE00000007, INF-ACT). This publication was produced while Silvia Fabi attending the PhD programme in Sustainable Development and Climate Change at the University School for Advanced Studies IUSS Pavia, Cycle XXXVIII, with the support of a scholarship financed by the Ministerial Decree no. 351 of 9th April 2022, based on the NRRP - funded by the European Union - NextGenerationEU - Mission 4 “Education and Research”, Component 1 “Enhancement of the offer of educational services: from nurseries to universities”; and while Mariachiara Vardeu was attending the PhD National Programme in One Health approaches to infectious diseases and life science research, Department of Public Health, Experimental and Forensic Medicine, University of Pavia, Pavia, 27100, Italy.

## DATA AVAILABILITY

All relevant data are contained within the manuscript.

## FIGURE LEGENDS

**Table 1.** Information on *Ixodes ricinus* samples positive for ALSV.

| Municipality | Collection date | Sample ID | Sample type | Ct | Isolate name / GenBank accession numbers |
| --- | --- | --- | --- | --- | --- |
| Belluno | March 09, 2021 | C19 | 10 nymphs | 27.46 | PX930134, PX930139, PX930175 |
|  |  | C20 | 10 nymphs | 28.23 | PX930141, PX930171 |
|  | March 17, 2021 | C53 | 10 nymphs | 36.00 | / |
|  |  | C55 | 10 nymphs | 27.50 | / |
|  | March 23, 2021 | C44 | 10 nymphs | 25.90 | PX930144, PX930162, PX930187 |
|  |  | C64 | 10 nymphs | 33.34 | / |
|  |  | C65 | 10 nymphs | 27.86 | PX930184 |
|  |  | C66 | 7 nymphs | 35.93 | / |
|  | April 20, 2021 | C97 | 10 nymphs | 23.88 | PX930129, PX930137, PX930163, PX930181 |
|  |  | C98 | 10 nymphs | 36.30 | / |
|  |  | C99 | 10 nymphs | 35.48 | / |
|  |  | C100 | 10 nymphs | 32.69 | PX930152, PX930172 |
|  |  | C101 | 10 nymphs | 35.26 | / |
| Sospirolo (Belluno) | March 03, 2021 | C6 | 10 nymphs | 36.92 | / |
|  | March 30, 2021 | C67 | 10 nymphs | 24.00 | PX930146, PX930160, PX930173 |
|  |  | C68 | 10 nymphs | 25.23 | PX930147, PX930180 |
|  |  | C73 | 10 nymphs | 26.85 | PX930138, PX930186 |
|  |  | C84 | 1 male | 20.36 | PX930133, PX930142, PX930167, PX930176 |
|  | May 21, 2021 | C215 | 10 nymphs | 25.94 | PX930128, PX930161, PX930182 |
|  | May 26, 2021 | C153 | 1 male | 37.53 | / |
|  |  | C225 | 10 nymphs | 27.40 | PX930145, PX930168, PX930183 |
|  | June 1, 2021 | C235 | 10 nymphs | 27.75 | PX930150, PX930164, PX930178 |
|  | June 11, 2021 | C284 | 10 nymphs | 38.00 | / |
|  | July 16, 2021 | C316 | 10 nymphs | 23.95 | PX930140, PX930166, PX930179 |
| Laghi-Posina (Vicenza) | May 12, 2022 | 21 | 10 nymphs | 35.93 | / |
|  |  | 24 | 10 nymphs | 37.44 | / |
|  |  | 39 | 1 male | 26.00 | PX930135, PX930188 |
|  |  | 42 | 1 male | 36.18 | / |

## Notes

### Competing Interest Statement

The authors have declared no competing interest.

### Summary of Updates

The revised version now includes the supplementary materials file associated with the article.

