## Supplementary material for "First detection and characterization of Alongshan virus in *Ixodes ricinus* ticks from Italy, 2021-2022": Supplemetary materials

**Appendix materials**

Table A1. Primers used for the amplification and sequencing of ALSV genomic segments

| **Name** | **Primer** | **5' Position*** | **GC-content %** | **Tm** |
| --- | --- | --- | --- | --- |
| seg1 FW1 | AGTTAATAGGAGCCAGCCTCA | 2 | 47.6 | 58.2 |
| seg1 RV1 | ACTCTTCCTTCCCTCTTTCCC | 594 | 52.4 | 58.1 |
| seg1 FW2 | TCGAAAGGATACGACAAGCTC | 423 | 47.6 | 57.9 |
| seg1 RV2 | GATCTGGGTTTCTAGCCTGTC | 947 | 52.4 | 57.8 |
| seg1 FW3 | GATACCGCCGAGAAACTTTGT | 773 | 47.6 | 58.7 |
| seg1 RV3 | GAGCACTCACGAATCTGTTCT | 1236 | 47.6 | 58.0 |
| seg1 FW4 | GACGGAGGAGGCTAAAGATAGA | 1052 | 52.4 | 57.9 |
| seg1 RV4 | CAGGCTCTGCTTAAAGGTGTT | 1589 | 47.6 | 58.5 |
| seg1 FW5 | AATAGGCGCGGAACCATGG | 1476 | 57.9 | 60.5 |
| seg1 RV5 | ATGAAACCTGTCCTCTGCCC | 2055 | 55.0 | 59.7 |
| seg1 FW6 | TGGTGGTCCAGAGAGAAATTG | 1987 | 47.6 | 57.6 |
| seg1 RV6 | ACTCCGGGATATGGTGGTAAT | 2550 | 47.6 | 58.0 |
| seg1 FW7 | CATTACATGCCGGTGAGAGAC | 2406 | 58.8 | 52.4 |
| seg1 RV7 | TTGCGCCGTGGCAAG | 3059 | 58.0 | 56.7 |
| seg2 FW1 | AAAGCGGACCCTTTCAGTTG | 1 | 50.0 | 58.7 |
| seg2 RV1 | GGTACAATGTAAGAGGTGGCC | 556 | 52.4 | 58.4 |
| seg2 FW2 | CTCCTCCCACTTATCGTCGC | 365 | 60.0 | 60.0 |
| seg2 RV2 | CCAGATGGTCATCTCAGAACG | 848 | 52.4 | 58.2 |
| seg2 FW3 | ACAGGACAAAGGCCATCTGG | 745 | 55.0 | 60.0 |
| seg2 RV3 | TCGGAGGAGATGATCATGGTT | 1213 | 47.6 | 58.3 |
| seg2 FW4 | AGGATCAGAGGTGACCCTCC | 995 | 60.0 | 60.0 |
| seg2 RV4 | CGATACCTCCCGTGATGTGG | 1523 | 60.0 | 60.0 |
| seg2 FW5 | GTCCATGAGGGCTTTTACCGAT | 1433 | 47.6 | 58.1 |
| seg2 RV5 | GTTGATGAGGAGAAGGTGAGC | 1873 | 52.4 | 58.4 |
| seg2 FW6 | CACTCTTGTTTCGTCACCGGA | 1833 | 55.0 | 60.0 |
| seg2 RV6 | CTCAGCCGGATAGAGGTAGAA | 2236 | 52.4 | 58.1 |
| seg2 FW7 | TTCAGTTGACTAYGTGACGGC | 2095 | 47.6 | 58.3 |
| seg2 RV7 | GACTCCGGTGGACCCC | 2763 | 75.0 | 58.1 |
| seg3 FW1 | TCCCGGGGGAGTTAATATGGA | 13 | 52.4 | 59.8 |
| seg3 RV1 | TGCTATGGACAGCATCACCA | 590 | 52.4 | 60.4 |
| seg3 FW2 | CGCTATAGTGGTGCTTACAGT | 506 | 47.6 | 57.6 |
| seg3 RV2 | CAGCTGTCTGACATACTCTGG | 1229 | 52.4 | 57.8 |
| seg3 FW3 | GACCATAGGCAGGAGCATAAG | 1055 | 52.4 | 57.9 |
| seg3 RV3 | GTACACGTGATCCACTATCGG | 1628 | 52.4 | 58.0 |
| seg3 FW4 | GAGGTCCGTGTGTCTATGGA | 1578 | 52.4 | 58.5 |
| seg3 RV4 | CGGTAGTACTCTCCCTCTCTT | 2032 | 52.4 | 57.4 |
| seg3 FW5 | AAGGAGTCATAACCCATCCA | 1948 | 52.4 | 58.1 |
| seg3 RV5 | TTGCAACGGGCATAGTGGAA | 2801 | 52.4 | 60.3 |
| seg4 FW1 | AACCAGCTCAGGCAGCAAGT | 12 | 57.9 | 59.6 |
| seg4 RV1 | CTGTAGCTCCATCCAAATTCCA | 521 | 45.5 | 58.1 |
| seg4 FW2 | CGGTGAGGAGATAGCACTCA | 330 | 55.0 | 58.6 |
| seg4 RV2 | TTCCCCTCCCCTACTGAAAAA | 887 | 47.6 | 58.3 |
| seg4 FW3 | AAGCTGGAAAGATCACCCAT | 800 | 45.0 | 56.8 |
| seg4 RV3 | TGATACTGTATCGTGGGCAG | 1495 | 50.0 | 56.8 |
| seg4 FW4 | ACGATATTCTTGGCGGCCAC | 1425 | 55.0 | 60.8 |
| seg4 RV4 | TCTGCTCATGGCCATGGTTAT | 1970 | 47.6 | 59.5 |
| seg4 FW5 | TCACGGACACGGGAGAGA | 1846 | 61.1 | 59.6 |
| seg4 RV5 | CTCACACATCAGTTTGCCCTG | 2452 | 55.0 | 58.8 |
| seg4 FW6 | AAGGTCCTGATGGTCCTGTC | 2358 | 55.0 | 58.7 |
| seg4 RV6 | CTTCCGGAACAAGAAACTAGC | 2720 | 47.6 | 56.9 |

*These coordinates refer to the Alongshan virus sequence with the NCBI reference codes MW094153, MW094159, MW094156, and MW094158 for sequence reference segments 1, 2, 3, and 4, respectively.

Table A2. Mutation rates were calculated for all Italian ALSV strains using the European reference sequence (Finland, 2011) as comparator. Rates were normalized to the number of effectively covered nucleotides, excluding gaps and ambiguous bases. Across all genomic segments, a consistent predominance of synonymous over non-synonymous substitutions was observed

| **Segment** | **Protein/CDS** | **Synonymous vs non-synonymous substitutions per 1,000 nt** | **Substitutions detected in all Italian isolates / conserved in most ALSV sequences** | **Lower-frequency substitutions in Italian strains** | **Figure** |
| --- | --- | --- | --- | --- | --- |
| 1 | NS5-like protein | 45.9 vs 4.9 | R185K, K530R, K744R, S860C | K553R (n=2); K346R, A563Y, V612S, D695E, T699K, Y858N (single strain) | S4 |
| 2 | VP1a, putative glycoprotein | 9.8 vs 3.7 | R46Q, Y424C, P477L | K311R (n=7); S443L (n=5); Q425R, S468L (n=2); P473L shared by 2 Italian sequences and 4 ALSV reference sequences; C26W, S144A, T216N, L273F, R346M, P357S, R475L (single strain) | S5 |
|  | VP1b, putative glycoprotein | 15.6 vs 1.5 | Not detected | L113F (n=4); R92K (n=3); N91K, V117A (n=2); V117A shared by 2 Italian sequences and reference PQ900136; L88P (single strain) | S5 |
| 3 | NS3-like protein | 55.0 vs 4.3 | I31V, R506K | V483L, Y489C (n=2); P57S, A123G, G182R, K241M, A266D, G308D, T330I, M364T, Y316H, H494L, T589S, V600G, E628G (single strain) | S6 |
| 4 | Capsid protein | 52.5 vs 2.2 | R156K, conserved across most available ALSV sequences, except for the Finnish reference strain | K55N (n=2); A173G (n=2); T63I, A83G, H170Y, H233L (single strain) | 2 |
|  | Membrane protein | 48.3 vs 5.1 | H250Q, V257E, I332V, T469A, K473R, conserved across most ALSV reference sequences, with divergence in the Finnish strain | L279F (n=2); K98R (n=1 Italian isolate; also detected in two Chinese strains); K299R, T264P, S272T and a few additional substitutions detected in single Italian strain | 2 |

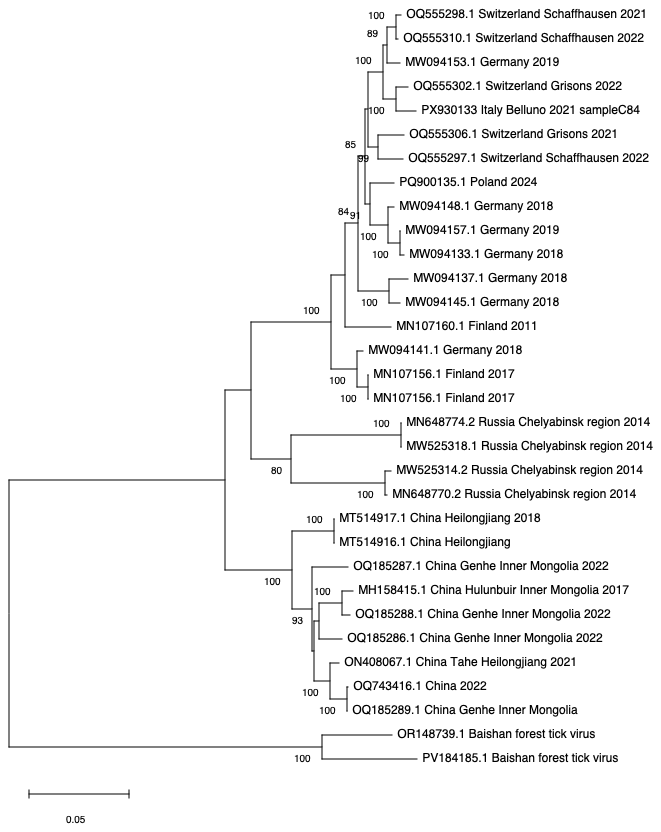

**Figure A1.** Phylogenetic tree of ALSV segment 1 sequences, including one strain detected in *Ixodes ricinus* ticks, northeastern Italy, 2021–2022. Italian sequences cluster within the European ALSV diversity. The tree was inferred using the Maximum Likelihood method with the Tamura–Nei model and generated using MEGA. Bootstrap values (>75%, 500 replicates) are shown. Scale bar indicates nucleotide substitutions per site.

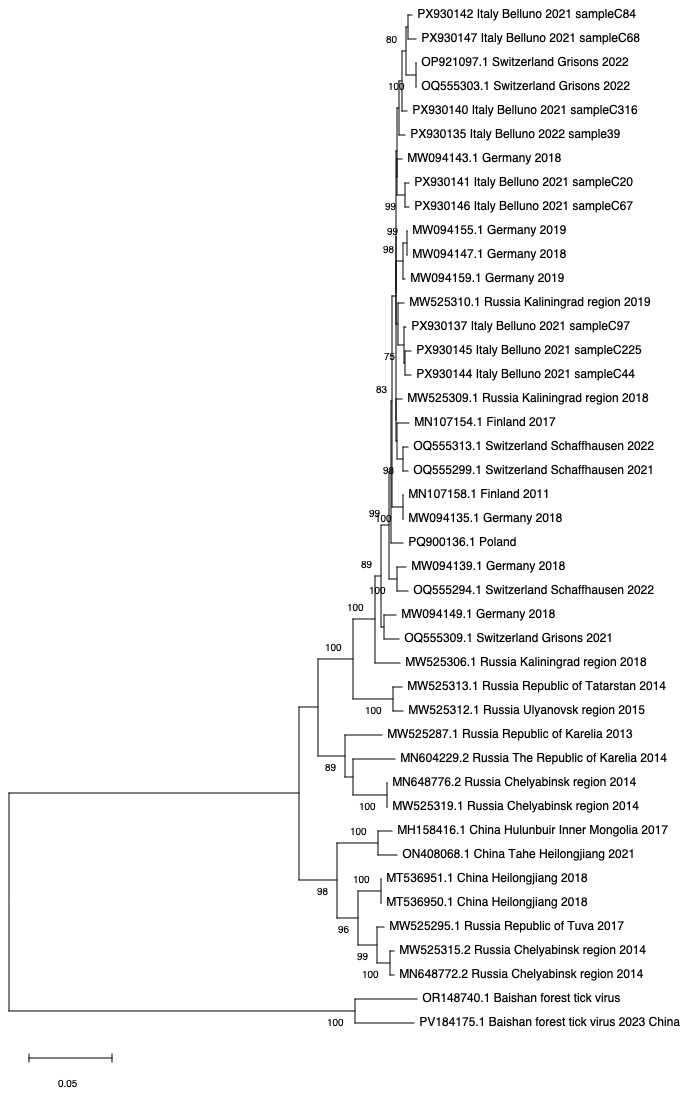

**Figure A2.** Phylogenetic tree of ALSV segment 2 sequences, including strains detected in *Ixodes ricinus* ticks, northeastern Italy, 2021–2022. Italian sequences cluster within the European ALSV diversity. The tree was inferred using the Maximum Likelihood method with the Tamura–Nei model and generated using MEGA. Bootstrap values (>75%, 500 replicates) are shown. Scale bar indicates nucleotide substitutions per site.

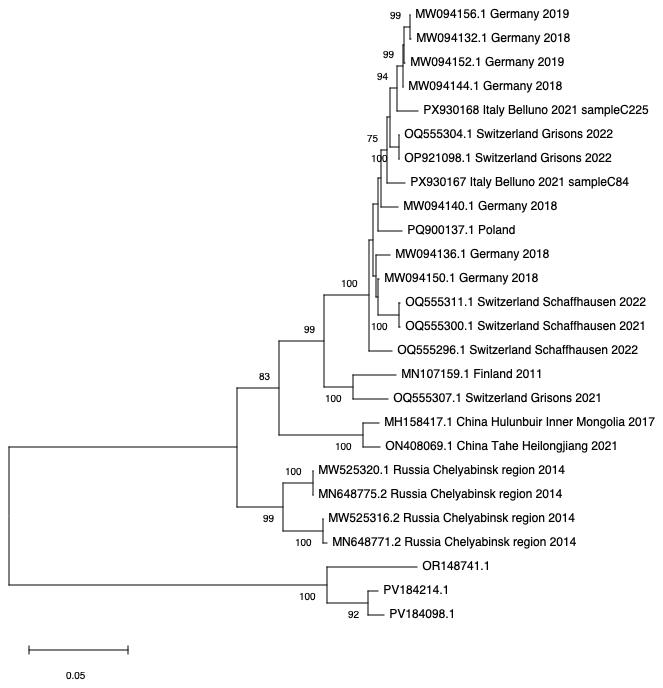

**Figure A3.** Phylogenetic tree of ALSV segment 3 sequences, including two strains detected in *Ixodes ricinus* ticks, northeastern Italy, 2021–2022. Italian sequences cluster within the European ALSV diversity. The tree was inferred using the Maximum Likelihood method with the Tamura–Nei model and generated using MEGA. Bootstrap values (>75%, 500 replicates) are shown. Scale bar indicates nucleotide substitutions per site.

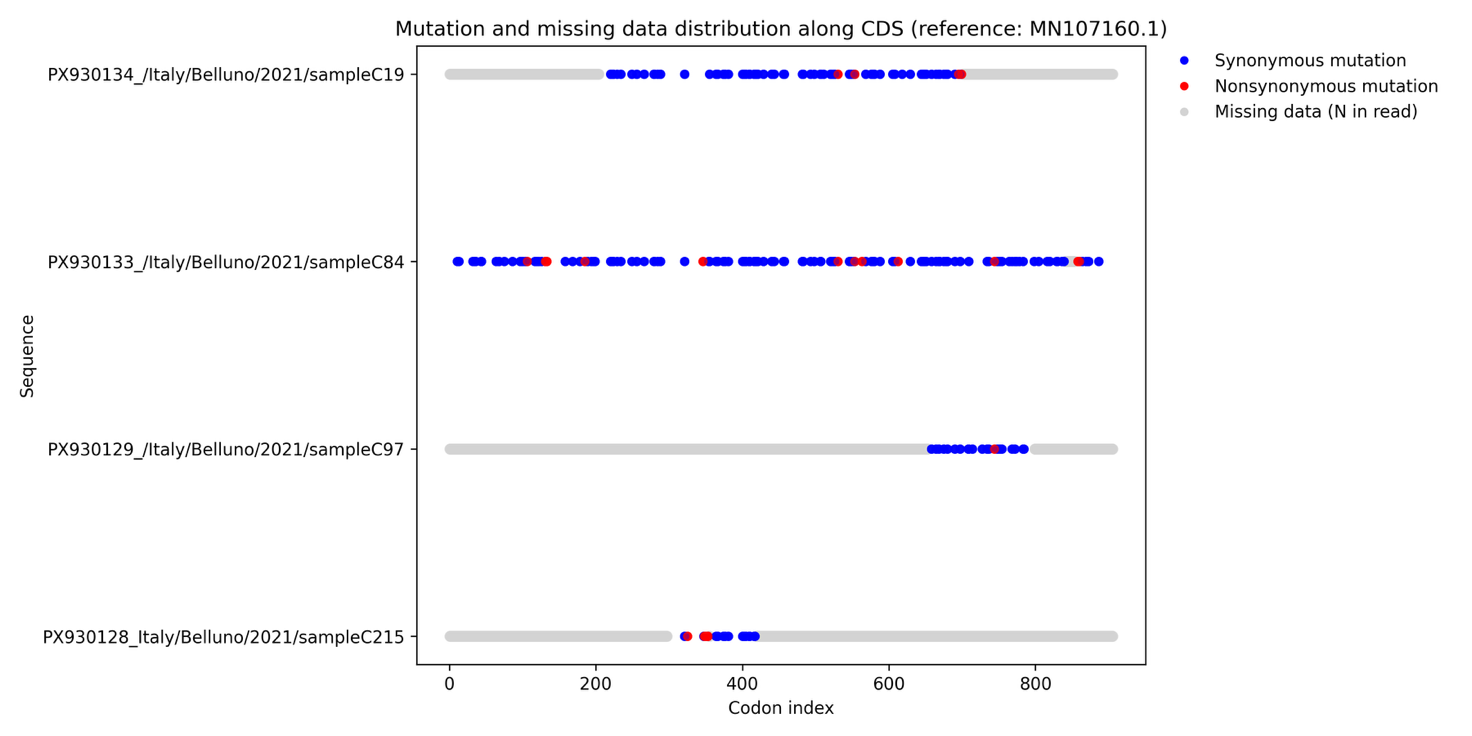

**Figure A4.** **Summary of all mutations detected in segment 1 of Italian ALSV strains relative to the Finnish European reference sequence.** Synonymous and non-synonymous mutations are reported as blue and red dots, respectively.

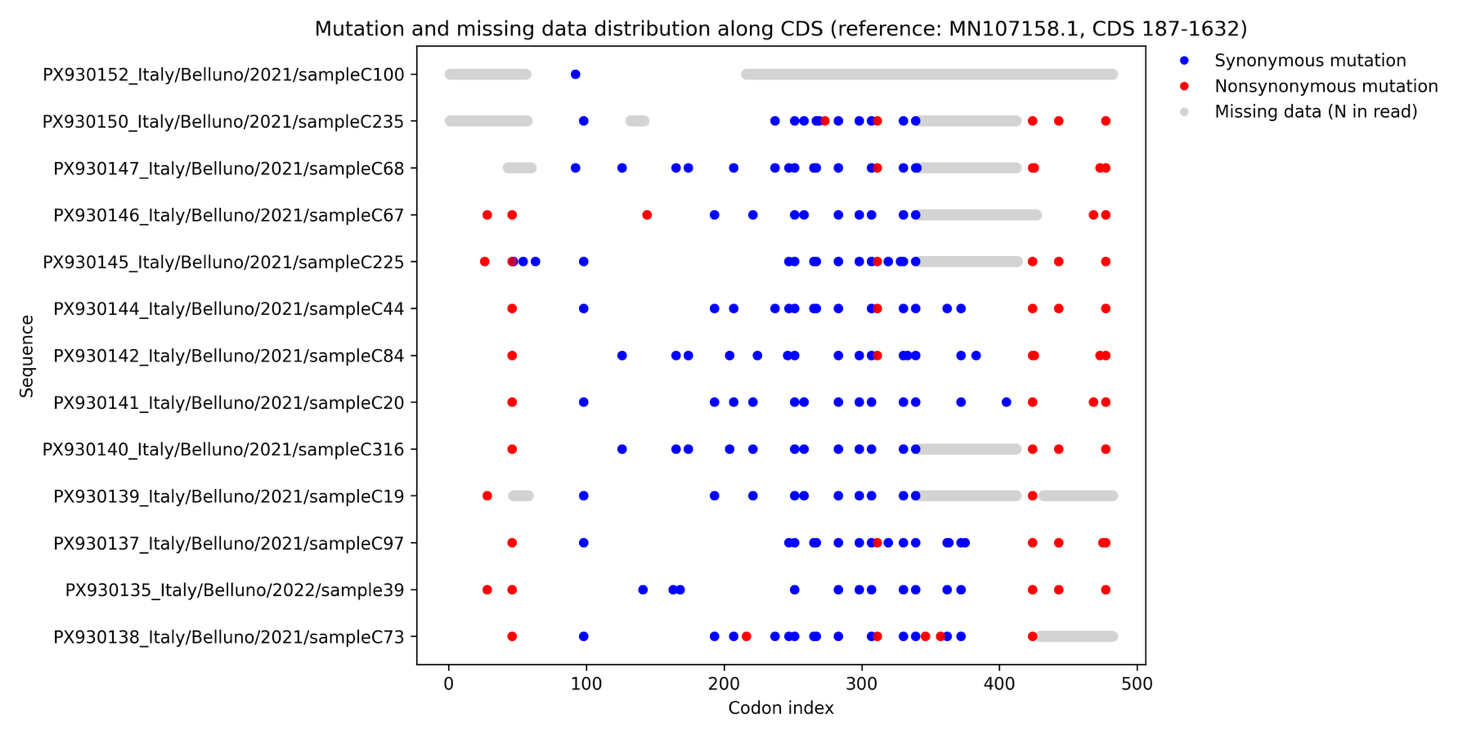

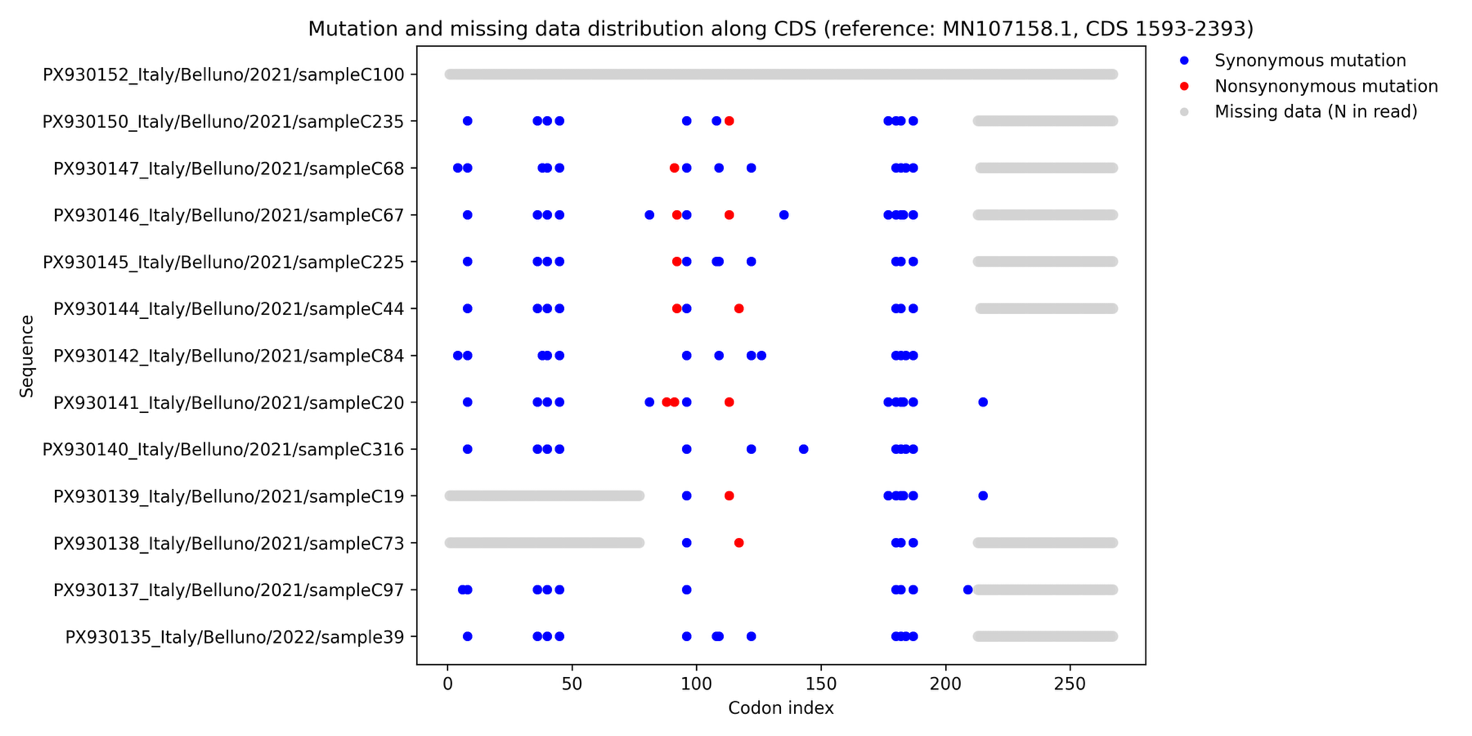

**Figure A5.** **Summary of all mutations detected in segment 2 of Italian ALSV strains relative to the Finnish European reference sequence.** Synonymous and non-synonymous mutations are reported as blue and red dots, respectively.

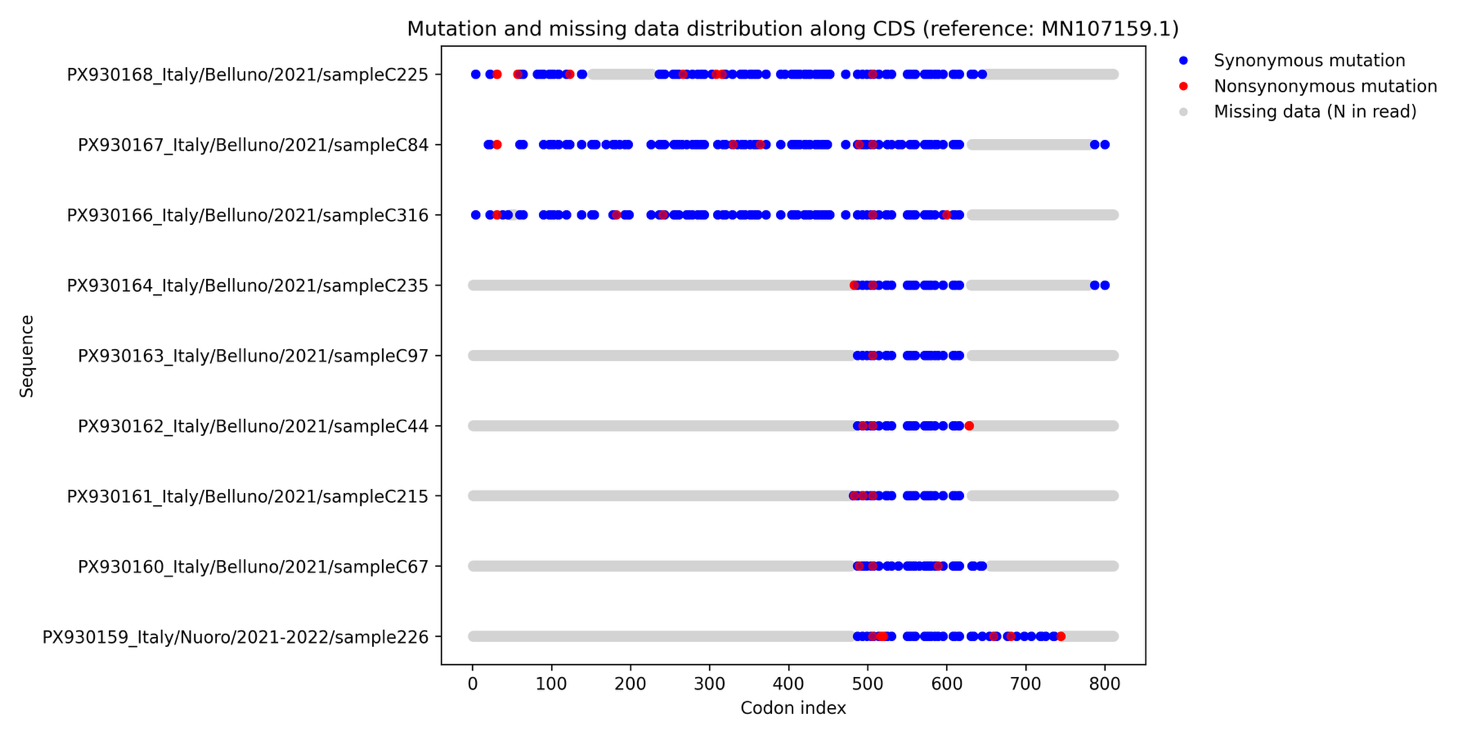

**Figure A6.** **Summary of all mutations detected in segment 3 of Italian ALSV strains relative to the Finnish European reference sequence.** Synonymous and non-synonymous mutations are reported as blue and red dots, respectively.
